# An unconstrained Euclidean parametrisation of phylogenetic tree space

**DOI:** 10.64898/2026.10.14.759122

**Authors:** Xavier Didelot

**Affiliations:** School of Life Sciences, University of Warwick, Coventry, United Kingdom; Department of Statistics, University of Warwick, Coventry, United Kingdom

**Keywords:** phylogenetic trees, Euclidean parametrisation, sampling tree distributions, Bayesian phylogenetic inference

## Abstract

Phylogenetic trees combine a discrete topology with continuous branch lengths, which makes them difficult to work with. We introduce a recursive contour-grafting construction that encodes both components using unconstrained real-valued coordinates. After fixing an order of the leaves, each new leaf is attached at a point selected by a contour traversal of the tree constructed so far. Moving a graft coordinate continuously along the contour enables changes of topology when internal branches contract to zero. For trees with *n* labelled leaves, we apply the construction to rooted ultrametric trees, rooted trees with unrestricted branch lengths, and unrooted trees, obtaining parametrisations of dimensions *n* − 1, 2*n* − 2, and 2*n* − 3, respectively. We show that the resulting construction functions are surjective and continuous onto the corresponding spaces of phylogenetic trees. We also determine the number of representations of a generic tree and calculate the Jacobian of each parametrisation. These results provide explicit change-of-variables formulae between Euclidean coordinates and the phylogenetic tree spaces, and we show how probability distributions on tree space can be sampled using generic algorithms that target unconstrained Euclidean coordinates. This approach paves the way for the development and application of new methodology in Bayesian phylogenetic inference.

## 1 Introduction

Phylogenetic trees are typically represented as the combination of a discrete topology and continuous branch lengths (Wiley and Lieberman 2011). This representation is natural, but it makes tree space difficult to explore. In particular, changing topology ordinarily requires purpose-built operations such as nearest-neighbour interchange (NNI) or subtree pruning and regrafting (SPR), whose performance can depend strongly on how exactly they are designed (Höhna and Drummond 2012; Zhang et al. 2020; Meyer 2021). It also means that many recent computational methods cannot be directly used in phylogenetics as they are designed to operate on vectors in a fixed Euclidean space.

A geometric description of tree space is provided by the Billera–Holmes–Vogtmann (BHV) space (Billera et al. 2001). Each tree topology is represented by an orthant of branch-length coordinates, and different orthants meet where one or more internal branches have length zero. This construction makes clear that topology changes can be viewed continuously: a branch belonging to one topology contracts before a branch belonging to another topology grows. The resulting space is nevertheless stratified, and computational methods that move between its orthants generally require special careful design (Dinh et al. 2017; Helekal et al. 2025). A few recent studies have sought to develop alternative continuous or vector-based representations of phylogenetic trees. For example, GradME uses continuous optimisation for tree inference under a balanced minimum-evolution criterion (Penn et al. 2023). Phylo2Vec presents a bijective integer-vector representation of binary tree topologies (Penn et al. 2025). These methods illustrate the potential of alternative representations for exploring tree space, but they address different objectives from the one considered here.

Here we develop a representation in which both topology and branch lengths are encoded by a single vector of unconstrained real parameters. The construction is recursive. After fixing an arbitrary order of the leaves, each new leaf is attached to the tree formed by the preceding leaves. Its graft point is selected by following a contour traversal of the current tree, so that continuous movement of a real-valued coordinate moves the graft point along branches and through the tree nodes. In this way, changes of topology arise without introducing a separate discrete topology variable. We show that this contour-grafting construction can be used to represent the three most widely used classes of phylogenetic trees, namely rooted ultrametric trees, rooted trees with unrestricted branch lengths and unrooted trees. Our main contribution is to characterise the theoretical properties of this parametrisation: we establish its continuity and surjectivity, determine the number of generic representations of each tree, and derive its Jacobian almost everywhere. These properties yield explicit change-of-variables formulae for transferring probability densities between tree space and Euclidean coordinates. The resulting framework provides a basis for applying algorithms defined on continuous parameter spaces to Bayesian phylogenetic inference, although the computational advantages of particular algorithms are beyond our scope.

The remainder of the article is organised as follows. Section 2 introduces the recursive contour-grafting construction. Sections 3–5 apply it to rooted ultrametric, rooted unrestricted, and unrooted trees, respectively. Section 6 establishes the link between probability densities in tree space and in the new parametrisations. Section 7 shows that the phylogenetic tree space can be effectively sampled via the new parametrisations, and Section 8 shows a small toy example of performing Bayesian phylogenetic inference. Section 9 describes details of how the methods were implemented and finally Section 10 discusses how the theory presented here could be used as a starting point for the development of new phylogenetic methods.

## 2 Recursive contour-grafting construction

All our parametrisations are constructed recursively using the same basic principle. We consider an arbitrary but fixed order for the leaves. The tree is constructed by adding each leaf one by one to the tree built so far. Let *T*__*i*__ denote the rooted tree formed by the first *i* leaves and let *L*(*T*__*i*__) be the sum of the branch lengths in the tree *T*__*i*__. The tree *T*__1__ consists only of the first leaf. When adding each leaf *i* beyond the first one, a graft point *x*__*i*__ is selected on or above *T*__*i*−1__ corresponding to the point where the new leaf is attached. This process is repeated for all leaves from the second one (*i* = 2) to the last one (*i* = *n*), each time adding to the tree *T*__*i*−1__ built previously.

The location of a graft point *x*__*i*__ is parametrised using a variable *a*__*i*__ ∈ R. If 0 ≤ *a*__*i*__ ≤ 2*L*(*T*__*i*−1__) then *a*__*i*__ corresponds to the distance traveled along the contour traversal of *T*__*i*−1__ before reaching the graft point. Starting at the root, the traversal follows each branch once away from the root and once back towards it, visiting the first child of each internal node and then the second. Let |*T* | denote the set of points on the tree *T*. The process of finding this graft point on a tree *T* using the contour traversal is denoted *γ*__*T*__ : [0, 2*L*(*T*)] → |*T* |.

We further define the extended contour traversal as illustrated in Figure 1 and described below. If *a*__*i*__ *>* 2*L*(*T*__*i*−1__), the grafting point is above the root of *T*__*i*−1__ by length *a*__*i*__ − 2*L*(*T*__*i*−1__). Likewise, if *a*__*i*__ *<* 0 then the grafting point is above the root of *T*__*i*−1__ by length −*a*__*i*__. This extended contour traversal is therefore a function Γ__*T*__ : ℝ → |*T* |^+^. The co-domain |*T* |^+^ is obtained by adding to |*T* | the points on a branch above the root at a distance *d* ∈ [0, ∞), with zero identified with the root. The extended contour traversal is defined as:

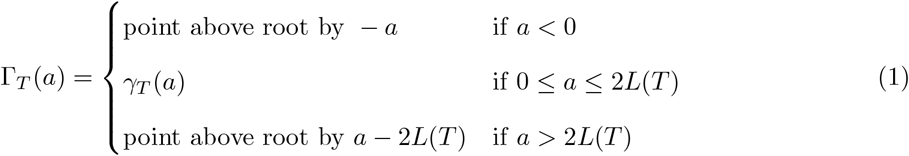

**Figure 1:**
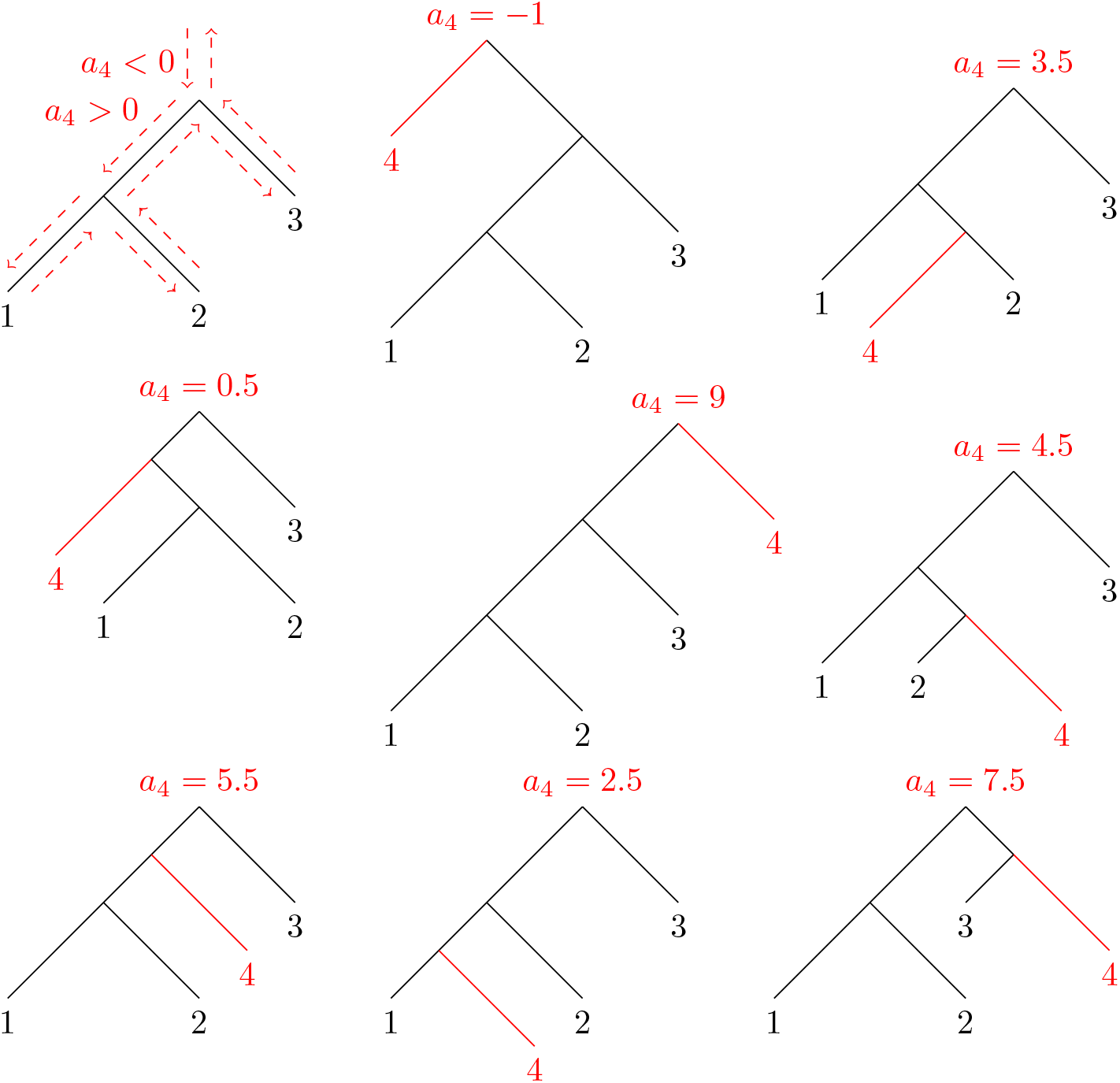
Illustration of the extended contour traversal. The initial tree is shown in black, with three leaves labelled 1, 2 and 3 and all four branches of length 1. Leaf 4 is added with the value of *a*_4_ determining the graft point as shown and the new terminal branch shown in red. When the contour traversal is traveling down (cases *a*_4_ = −1, *a*_4_ = 3.5 and *a*_4_ = 0.5) then the leaf becomes the first child and otherwise (remaining cases) the leaf becomes the second child of the newly created node at the graft point.

At the graft point *x*__*i*__ = Γ__*T*_*i−*1_ (*a*__*i*__), a new node is created with first child equal to what was previously below that point, and second child equal to the new branch of length *l*__*i*__ leading to the leaf *i*. Let *∆*__*T*__ (*a*) = 1 if the contour traversal was traveling down a branch when it reached the graft point *x*__*i*__, and *∆*__*T*__ (*a*) = 0 otherwise. If *∆*__*T*__ (*a*) = 1 then the two children of the new node are swapped. This convention also defines the ordering of the children used in subsequent contour traversals. It inserts the new leaf into the contour exactly where the graft point was reached: if the traversal is going down it meets the new node before the old subtree, and if it is going up it has just left the old subtree. The order in which the contour visits the subtrees around each node is therefore preserved when the graft point crosses that node, which is important to obtain continuity as we will see later. If *x*__*i*__ lies above the current root, the branch joining the old root to *x*__*i*__ is added to the tree and *x*__*i*__ becomes the new root. Above the root the direction of travel is defined as *∆*__*T*__ (*a*) = 1 for *a <* 0 and *∆*__*T*__ (*a*) = 0 for *a >* 2*L*(*T*). If *x*__*i*__ coincides with an existing node, the operation is interpreted as the corresponding limiting construction with a zero-length branch. Thus the construction takes values in the completion of binary tree space, while almost every parameter vector produces a binary tree with strictly positive branch lengths.

The extended contour traversal is continuous because *γ*__*T*__ (0) = *γ*__*T*__ (2(*L*(*T*))) = root(*T*). Every point in the interior of a branch of *T* is visited exactly twice, once in each traversal direction. A point at distance *d >* 0 above the root also has exactly two preimages, namely −*d* and 2*L*(*T*) + *d*. Tree nodes are exceptional because they may be visited more than twice; these exceptional points form the boundaries between contour regions. By considering the three cases (Equation 1) in which leaves can be added to the tree and the effect each of them has on the branch lengths we can deduce the following recursive formula for the sum of the branch lengths *L*(*T*__*i*__):

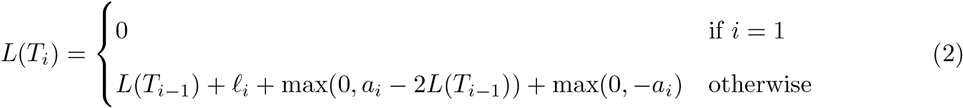

## 3 Rooted ultrametric trees

We first consider phylogenetic trees with *n* labelled leaves that are binary, rooted, and with the same distance from the root to each leaf (this property is called ultrametricity). We use the superscript U, standing for ultrametric, to denote the mathematical quantities relating to this type of tree. There are *n* − 1 internal nodes including the root. The topology of the tree is discrete and there are (2*n* − 3)!! = (2*n* − 2)!*/*(2^*n*−1^(*n* − 1)!) unique possibilities (Felsenstein 1978). The branch lengths are continuous but not independent from each other. Since the tree is ultrametric, the branch lengths are constrained so that the number of free parameters corresponds to the *n* − 1 values for the heights of the internal nodes relative to the contemporaneous leaves.

The previous section introduced the recursive principle for constructing trees using the extended contour traversal for grafting, with parameters (*a*__*i*__)__*i*=2,…,*n*__ ∈ ℝ^*n*−1^ used in Equation 1. To produce an ultrametric rooted tree, once a graft point *x*__*i*__ has been selected, the length *l*__*i*__ of the newly created terminal branch must be equal to the height of *x*__*i*__ relative to the existing leaves. Specifically, for a point *x* on an ultrametric tree *T*, let *h*__*T*__ (*x*) denote its distance to any descendant leaf. For a point at distance *d* above the current root *r*, its distance is *h*__*T*__ (*x*) = *h*__*T*__ (*r*) + *d*. The length of the new branch is simply *l*__*i*__ = *h*__*T*_*i−*1_ (*x*__*i*__). This construction is detailed in Algorithm 1. The dimensionality of the parameters is *n* − 1 which is the dimensionality needed to specify the *n* − 1 heights of the internal nodes. A small example of applying Algorithm 1 is shown in Figure 2.

**Figure 2:**
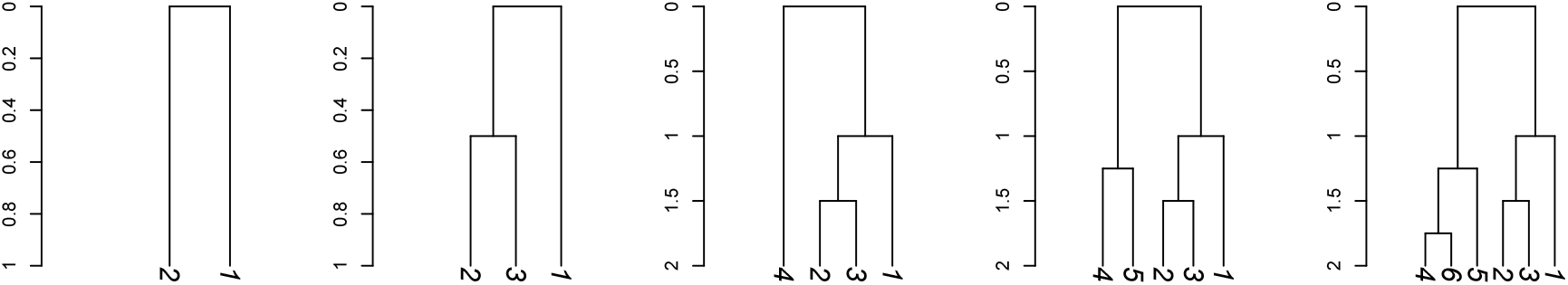
An example of construction of an ultrametric tree using as input *n* = 6 and (*a*_*i*_)_*i*=2,…,*n*_ = (−1, 1.5, −1, 2.75, 2.25).

### Algorithm 1

Construction of ultrametric rooted trees

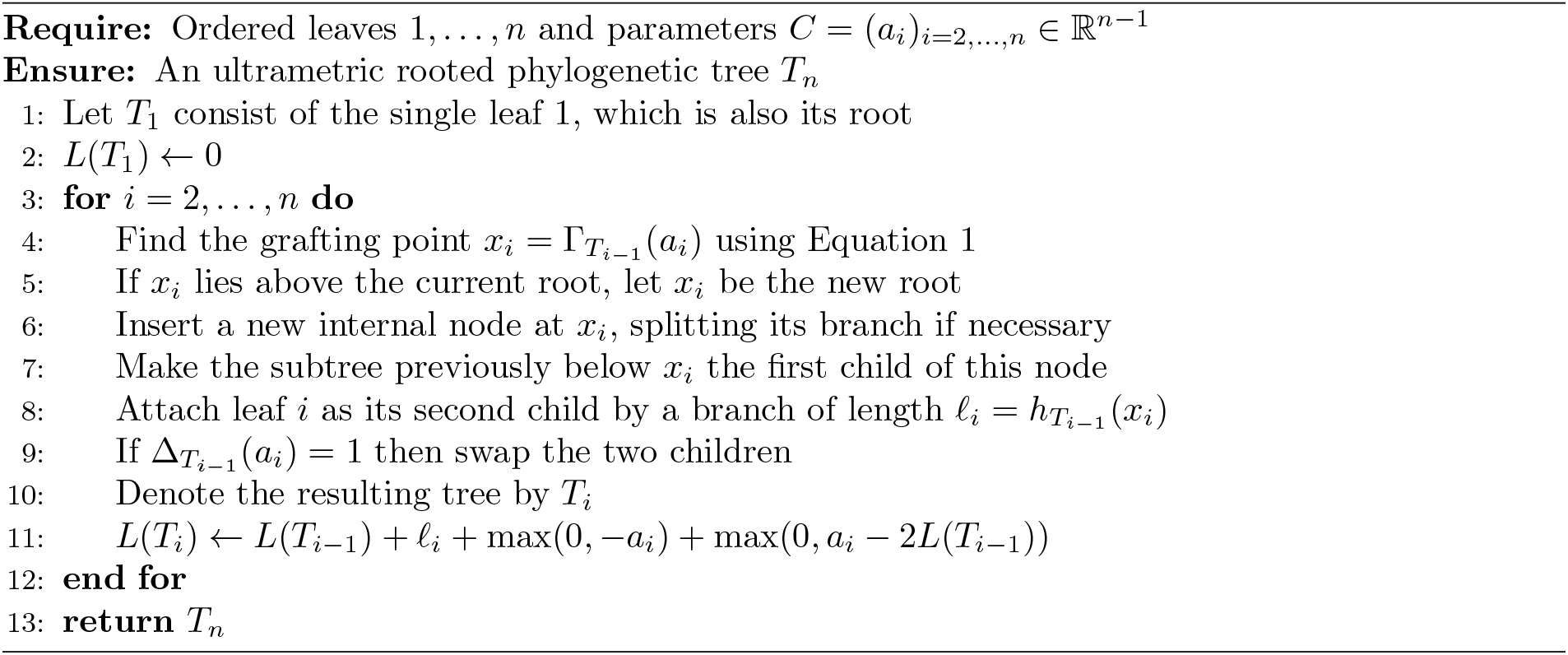

Let 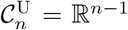 denote the parameter space. Let 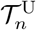 denote the space of rooted binary ultrametric trees with *n* labelled leaves and strictly positive branch lengths. Let 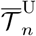 denote its completion obtained by allowing zero-length branches and identifying trees that define the same labelled metric tree after contraction of all zero-length branches. An element of 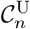 is the vector *C* = (*a*__*i*__)__*i*=2,…,*n*__, and the construction results in a unique tree in 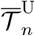. The construction in Algorithm 1 is therefore a function 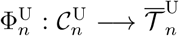. Almost every value of *C* produces a tree in 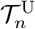; the exceptional values correspond to grafting at tree nodes, including the contour endpoints.

The function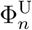is surjective onto 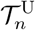, meaning that for any phylogenetic tree 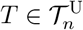 there is at least one parametrisation *C* such that 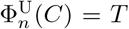. To see this, consider any 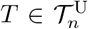, remove leaf *n* and its terminal branch, and suppress its resulting degree-two parent. Repeating this operation produces trees with successively fewer leaves. Reversing these steps reconstructs *T* using the proposed grafting procedure, because every required grafting location is reached by the extended contour traversal and ultrametricity uniquely determines the length of each newly added terminal branch. However, the function 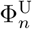 is not injective as there are more than one *C* such that 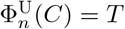. This is a consequence of each grafting point being visited twice by the contour traversal: once as the algorithm goes down the branch and then again as it goes back up the branch. Likewise, a grafting point above the root by length *x* can be obtained by both *a*__*i*__ = −*x* and *a*__*i*__ = 2*L*(*T*__*i*−1__) + *x*. The two representations of a grafting point give the two possible orders of the children of the new node, and this order determines the later contour traversals. It is therefore convenient to count the representations of a tree through the order of the children. A vector *C* determines *T* together with an order of the children at each of its *n* − 1 internal nodes, and *C* can be recovered from this ordered tree by removing leaves *n, n* − 1, …, 2 in turn: *a*__*i*__ is the position of the graft point on the contour of the remaining tree, at the descending visit if leaf *i* is the first child of its parent and at the ascending visit if it is the second child (this gives −*x* and 2*L*(*T*__*i*−1__) + *x* respectively above the root). Every order of the children of *T* arises in this way. For an ultrametric rooted binary tree *T* with strictly positive branch lengths, there are therefore 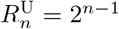 distinct values of *C* such that 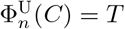. We call this number the redundancy of function 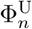. This duplication of grafting coordinates is needed to achieve continuity: it allows the grafting location to move continuously through the endpoints of the contour and across changes in tree topology, without an explicit jump between discrete topological states.

We now show that 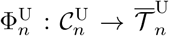 is continuous. Since the contour traversal depends on the order of the children, we consider trees whose children are ordered. Equip 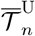 with the restriction of the completed rooted Billera–Holmes–Vogtmann metric (Billera et al. 2001), including terminal branch lengths. Within each region of parameter space in which the selected contour branches and traversal directions are fixed, all node heights and branch lengths vary continuously with the parameters. When a grafting location passes through an internal node, the separating internal branch contracts continuously to zero before the adjacent topology is entered. The two topologies therefore meet at the same boundary tree in 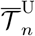. Both sides also give the same order of the children: the contour visits an internal node once for each pair of neighbouring subtrees (and once on each side of the branch above it), and the convention on *∆*__*T*__ inserts the new leaf at the corresponding place in the contour, whichever side the graft point is approached from. Subsequent contour traversals therefore also agree. When the grafting location passes through a leaf, the relevant terminal branch tends continuously to zero and the two contour directions likewise meet at the same boundary tree. The two directions give the two orders of the new leaf and the old leaf, but the terminal branch of the old leaf has length tending to zero, so the contour traversals of the two trees coincide. Similarly, the branch added above the root tends continuously to zero as *a*__*i*__ approaches either endpoint of the contour, namely at *a*__*i*__ = 0 or *a*__*i*__ = 2*L*(*T*__*i*−1__), and the order of the children is again the same on both sides (the new leaf is the first child at *a*__*i*__ = 0 and the second child at *a*__*i*__ = 2*L*(*T*__*i*−1__)). The graft location Γ__*T*__ (*a*) is itself jointly continuous in *a* and in the branch lengths of *T*, including when branches shrink to zero, because the contour is a closed walk of total length 2*L*(*T*) that depends only on the ordered tree. Since *T*__*i*−1__ varies continuously with (*a*__2__, …, *a*__*i*−1__), the continuity of each grafting step therefore carries over to the next, and applying this argument inductively establishes the continuity of 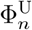. Within each binary topology orthant, the heights of the internal nodes provide local coordinates. Densities on tree space are understood with respect to counting measure on topologies and Lebesgue measure in these local height coordinates.

Finally we calculate the Jacobian of 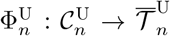 within a binary topology. Let ***h*** = (*h*, …, *h*) where *h*__*i*__ is the height of the internal node created when leaf *i* is grafted. Within a fixed contour region, 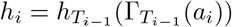 The new height depends only on *a*__2__, …, *a*__*i*__, not on later parameters. Moreover,

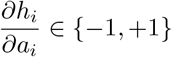

because height changes at unit rate along every fixed contour branch and along the branch above the root. Therefore the Jacobian matrix is triangular, with diagonal entries +1 or −1. Since the determinant of a triangular matrix is the product of the terms on its diagonal we have that the absolute Jacobian determinant is simply equal to:

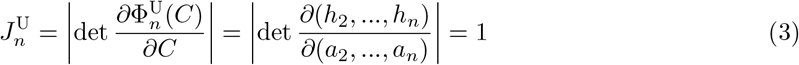

This identity holds away from contour-region boundaries, which form a set of Lebesgue measure zero in 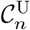.

## 4 Rooted trees with unrestricted branch lengths

We now consider phylogenetic trees that are binary and rooted, with *n* labelled leaves. We use the superscript R, standing for rooted, to denote the mathematical quantities relating to this type of tree. There are *n* − 1 internal nodes including the root. The topology of the tree is discrete and there are (2*n* − 3)!! = (2*n* − 2)!*/*(2^*n*−1^(*n* − 1)!) unique possibilities (Felsenstein 1978). The branch lengths are continuous, and there are 2*n* − 2 branch lengths taking values in ℝ_*>*0_ (one for each leaf and non-root internal node). Our construction is similar to the one above for ultrametric trees. The main difference is that there is now another parameter *b*_*i*_ ∈ ℝ for each leaf beyond the first one, that represents the length *l*_*i*_ = exp(*b*_*i*_) of the branch separating the graft point (determined by *a*_*i*_) and the newly added leaf *i*. This construction is detailed in Algorithm 2 and a small example of application is shown in Figure 3.

**Figure 3:**
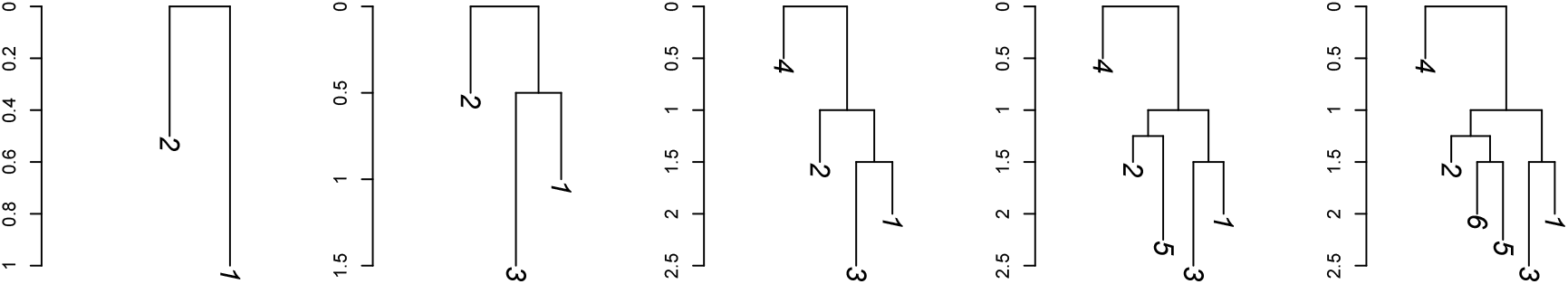
An example of construction of a rooted tree using as input *n* = 6, (*a*_*i*_)_*i*=2,…,*n*_ = (−1, 1.5, −1, 2.75, 3) and (*b*_*i*_)_*i*=2,…,*n*_ = log(0.5, 1, 0.5, 1, 0.5).

### Algorithm 2

Construction of rooted phylogenetic trees

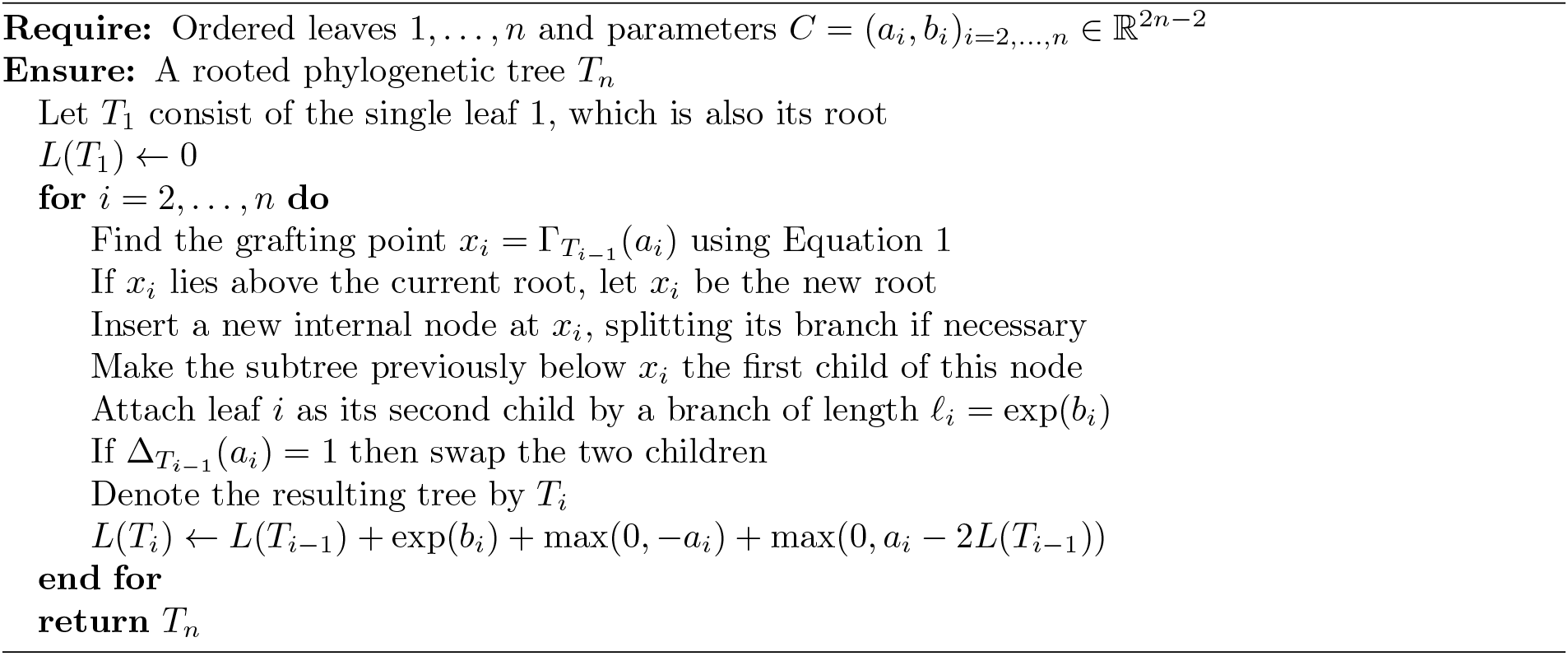

Let 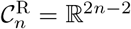 denote the parameter space. Let 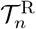 denote the space of rooted binary phylogenetic trees with *n* labelled leaves and strictly positive branch lengths. Let 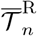 denote its completion obtained by allowing zero-length branches and identifying trees that define the same labelled metric tree after contraction of all zero-length branches. An element of 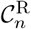 is the vector *C* = (*a*_*i*_, *b*_*i*_)_*i*=2,…,*n*_, and Algorithm 2 defines a function 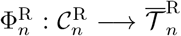. By the same arguments as in the ultrametric case, this function is continuous and is surjective onto 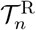. Almost every parameter vector produces a tree in 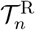, and every tree in 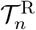 has exactly 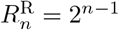 preimages, which we call the redundancy of 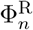.

As previously, we calculate the Jacobian of 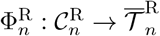 within a given binary topology. We consider this Jacobian locally, within a region of 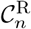where the resulting topology, the selected contour branch for each grafting point, and its traversal direction are fixed. Let ***l***^(*i*−1)^ denote the vector of branch lengths in *T*_*i*−1_. If leaf *i* is grafted in the interior of a branch of length *l*_*j*_, that branch is replaced by two branches of lengths

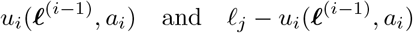

and a terminal branch of length exp(*b*_*i*_) is added. Along any fixed branch and direction of the contour traversal, *u*_*i*_ is affine in *a*_*i*_, with

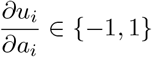

After reordering the branch-length coordinates, the absolute Jacobian determinant of this grafting step is therefore

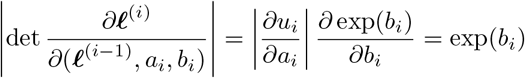

With a suitable ordering of the branch-length coordinates, the derivative of the full recursive construction is block triangular. Consequently, the determinants contributed by successive grafting steps multiply. The same result holds when the grafting point lies above the root, since the new root branch has derivative −1 or +1 with respect to *a*_*i*_. Applying recursively gives, for every *C* in which the grafting points do not coincide with a node or an endpoint of the contour:

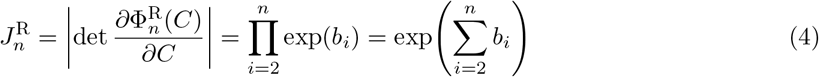

This identity holds away from contour-region boundaries, which form a set of Lebesgue measure zero in 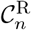.

## 5 Unrooted trees

Let us now consider unrooted binary phylogenetic trees with *n ≥* 3 labelled leaves. We use the superscript N, standing for not-rooted, to denote the mathematical quantities relating to this type of tree. Such a tree has *n*−2 internal nodes, 2*n*−3 branch lengths and (2*n*−5)!! = (2*n*−5)!*/*(2^*n*−3^(*n*−3)!) possible topologies (Felsenstein 1978). Briefly, we obtain an unconstrained parametrisation of dimension 2*n* − 3 by first applying the rooted construction of the previous section to the first *n* − 1 leaves to produce *T*_*n*−1_, and then grafting the last leaf *n* always at the temporary root of *T*_*n*−1_, after which the root is forgotten.

Let 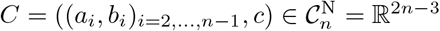. First construct a rooted tree *T*_*n*−1_ on leaves 1, …, *n* − 1 using Algorithm 2, with leaf 1 as the initial leaf and parameters (*a*_*i*_, *b*_*i*_)_*i*=2,…,*n*−1_. Then attach leaf *n* directly to the root of *T*_*n*−1_ by a branch of length exp(*c*), and then forget the root designation. The former root has degree three and therefore becomes an ordinary internal node of the resulting unrooted tree. This construction is detailed in Algorithm 3 and a small example of application is shown in Figure 4.

**Figure 4:**
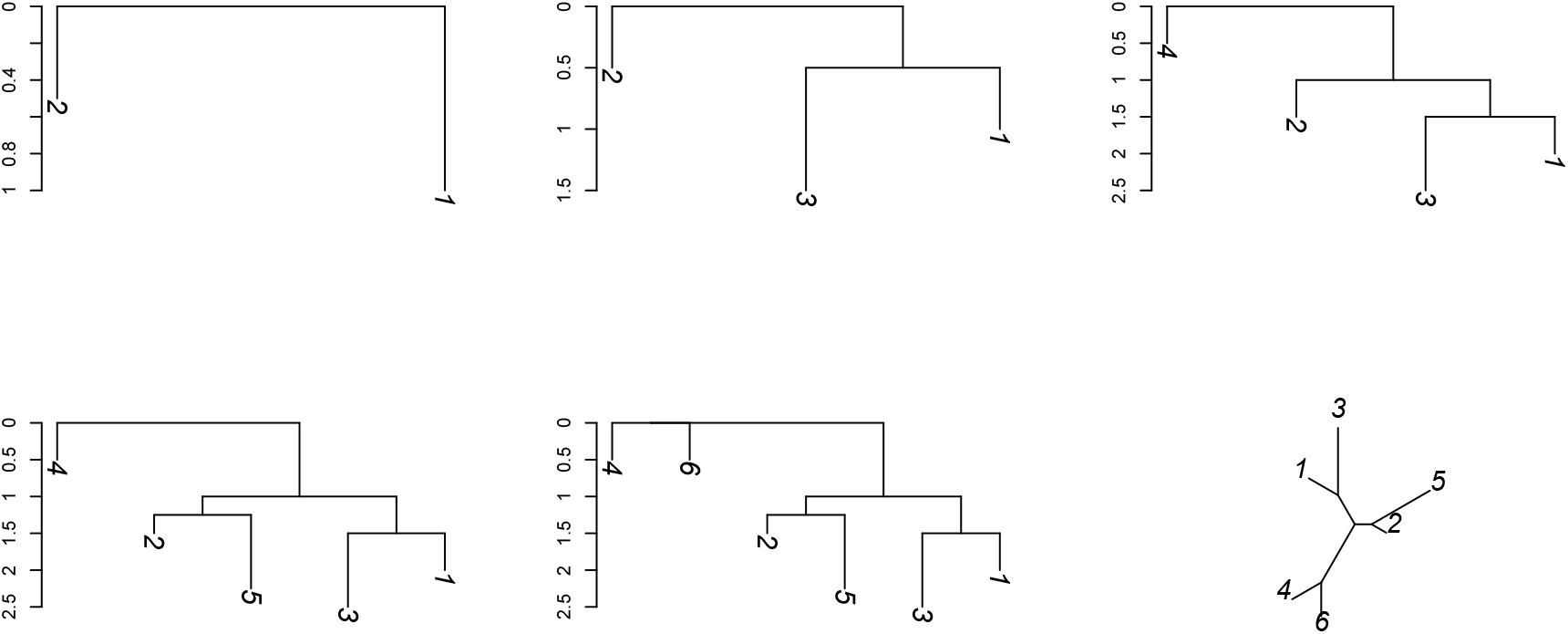
An example of construction of an unrooted tree using as input *n* = 6, (*a*_*i*_)_*i*=2,…,*n*−1_ = (−1, 1.5, −1, 2.75), (*b*_*i*_)_*i*=2,…,*n*−1_ = log(0.5, 1, 0.5, 1) and *c* = log(0.5).

Let 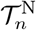 denote the space of unrooted binary phylogenetic trees with *n* labelled leaves and strictly positive branch lengths, and let 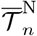 denote its completion obtained by allowing zero-length branches and identifying trees that define the same labelled metric tree after contraction of all zero-length branches. Algorithm 3 defines a function 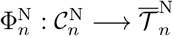. Almost every parameter vector produces a tree in 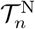; the exceptional values correspond to grafting at tree nodes, including contour endpoints.

### Algorithm 3

Parametrisation of unrooted phylogenetic trees

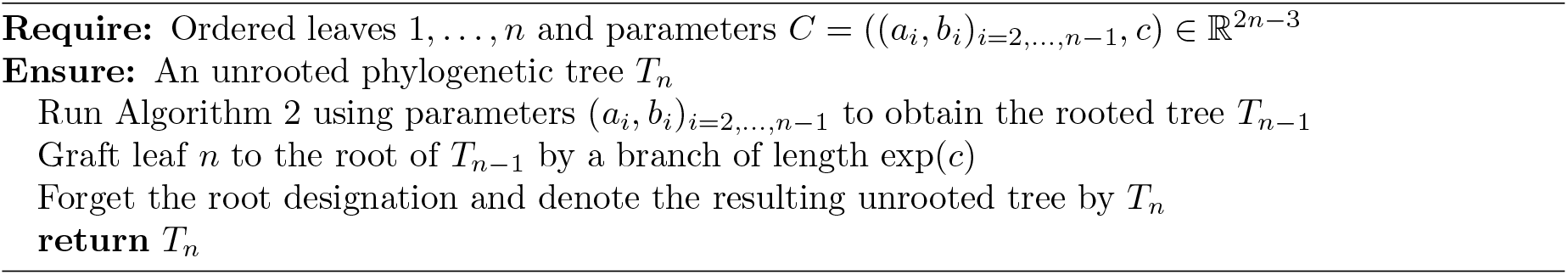

The function 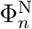 is surjective onto 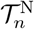. To see this, consider any 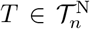. Remove leaf *n* and its terminal branch, and regard the resulting degree-two node as the root of the remaining tree on leaves 1, …, *n* − 1. This is a rooted binary phylogenetic tree and can therefore be represented by the function 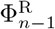 from the previous section. Setting *c* to the logarithm of the removed terminal branch length reconstructs *T*. Every tree in 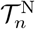 has exactly 2^*n*−2^ preimages. Indeed, the rooted tree on leaves 1, …, *n* − 1 has 2^*n*−2^ representations by the result of Section 4, while the length of the branch leading to leaf *n* uniquely determines *c*. Thus the redundancy of 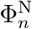 is 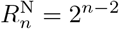

The function 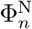 is continuous. Equip 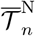with the completed unrooted Billera–Holmes–Vogtmann metric, including terminal branch lengths. The continuity of 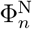 follows from the continuity of the function 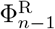, together with the continuity of attaching a terminal branch of length exp(*c*) at the root and then forgetting the root designation. At topology boundaries, zero-length branches are identified in 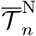 in the same way as in the rooted construction.

Finally, we calculate the Jacobian within a given binary topology. Use the 2*n* − 3 unrooted branch lengths as local coordinates. The rooted construction on leaves 1, …, *n* − 1 contributes exp ^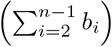^ to the absolute Jacobian (cf Equation 4), and the additional terminal branch has length exp(*c*) and derivative exp(*c*). With a suitable ordering of the coordinates, the full derivative is block triangular, and hence

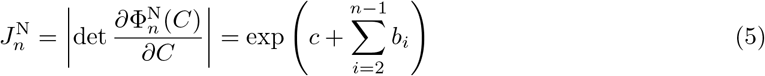

almost everywhere.

## 6 Distribution for each tree type

The main properties of the three constructions for ultrametric trees, rooted trees and unrooted trees are summarised and compared in Table 1. We now specify a probability model for each tree type. For ultrametric trees, we use the coalescent model (Kingman 1982a,b) which is the backward-in-time genealogical process for a sample of size *n* taken from large populations evolving under many forward-in-time models, such as the Wright-Fisher model (Wright 1931; Fisher 1930). Under this model, any pair of individuals finds a common ancestor at rate 1*/θ* where *θ* is a scaling factor on branch lengths. The model starts with the *n* individuals at time zero and builds the tree backward-in-time, so that the tree is ultrametric. Initially there are *n* individuals which means 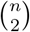 pairs and therefore the height of the first node is Exponentially distributed with rate 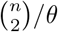 Any of the 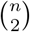 pairs is then equally likely to coalesce together, leaving *n* − 1 ancestors. By repeating this process and noting *τ*_*k*_ for the duration of the time interval during which there are *k* ancestors, the density of a tree is obtained as:

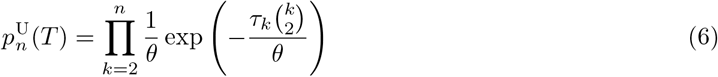

**Table 1:** Summary of the three parametrisations.

| Tree space | Parameters | Dimensions | Redundancy | Absolute Jacobian |
| --- | --- | --- | --- | --- |
| Rooted ultrametric (U) | $(a_i)_{i=2,\dots,n}$ | $n - 1$ | $2^{n-1}$ | 1 |
| Rooted unrestricted (R) | $(a_i, b_i)_{i=2,\dots,n}$ | $2n - 2$ | $2^{n-1}$ | $\exp\left(\sum_{i=2}^n b_i\right)$ |
| Unrooted (N) | $((a_i, b_i)_{i=2,\dots,n-1}, c)$ | $2n - 3$ | $2^{n-2}$ | $\exp\left(c + \sum_{i=2}^{n-1} b_i\right)$ |

Under the coalescent model, the distribution of the sum of branch lengths is (Tavaré 1984):

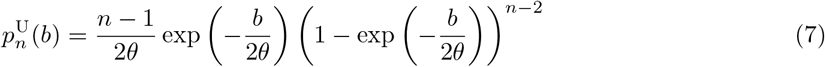

The distribution of the length of a single terminal branch is (Caliebe et al. 2007):

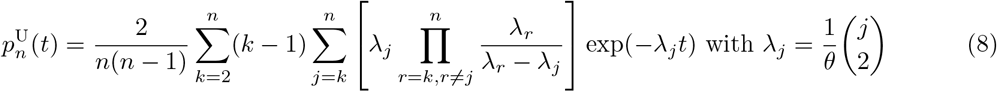

For rooted trees we consider a model in which all (2*n* − 3)!! topologies are equally likely and each of the 2*n* − 2 branch lengths *l*_*i*_ follow an Exponential distribution with mean *θ*, so that:

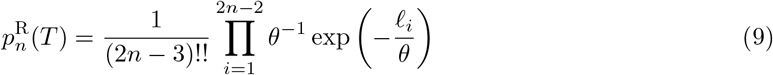

Under this model a terminal branch length and the sum of branch lengths are distributed as Exponential and Gamma, respectively:

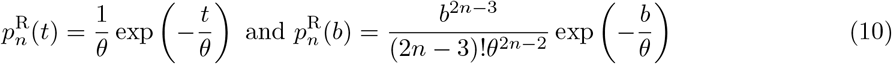

For unrooted trees we consider a model in which all (2*n* − 5)!! topologies are equally likely and each of the 2*n* − 3 branch lengths *l*_*i*_ follow an Exponential distribution with mean *θ*, so that:

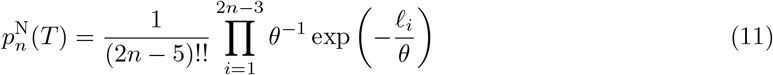

Under this model a terminal branch length and the sum of branch lengths are distributed as Exponential and Gamma, respectively:

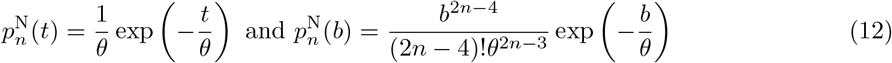

## 7 Sampling the tree distributions

Having equipped each of the three phylogenetic tree spaces with a distribution (Equations 6, 9 and 11), we now show how to sample from these distributions using our new Euclidean parametrisations. This requires using a change-of-variables formula that has the same form in all three cases. Let *j* ∈ {U, R, N} denote the type of tree under study. Let 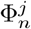 denote one of the three functions (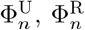 or 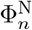), let 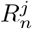 be its redundancy (cf Table 1), and let 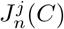 be its absolute Jacobian from Equations 3, 4 or 5. If 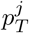 is a density on the corresponding tree space, with respect to counting measure on topologies and Lebesgue measure in the stated local coordinates, then assigning equal mass to all equivalent representations leads (almost everywhere) to the density:

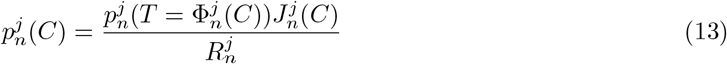

Thus the parametrisations provide a similar link between a density 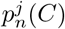 in unconstrained Euclidean space and a density 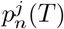 in phylogenetic space, for each of the three types of trees.

To sample from the distributions in Equation 13 we use an adaptive Metropolis sampler (Haario et al. 2001; Vihola 2011), specifically the robust adaptive Metropolis algorithm with coerced acceptance rate (Vihola 2012). This algorithm uses a multivariate normal distribution as the proposal kernel *q*(*C*^′^|*C*) which is symmetric (meaning that *q*(*C*^′^|*C*) = *q*(*C*|*C*^′^)), so that the acceptance probability is simply min(1, *p*_*C*_(*C*^′^)*/p*_*C*_(*C*)) calculated using Equation 13. The mean of the proposal kernel is the current value *C* and its covariance matrix is tuned to achieve an acceptance rate of 0.234. This target was derived for smooth high-dimensional targets which is not the case here, but should nevertheless be reasonable (Roberts et al. 1997; Andrieu and Thoms 2008). Note that this sampler by definition only operates on continuous parameters and could therefore not be applied directly to the phylogenetic tree spaces as usually defined. For each of the three models we take *n* = 5 to generate a small example, with *θ* = 0.01 as the scale of the branch lengths. In each case the sampler is run for 2 × 10^5^ iterations, with the first half discarded as burn-in and the remainder sampled every 10 iterations to produce a sample of size 10,000.

Under the coalescent model (Equation 6), the topologies are not equally likely but the ranked topologies (also known as histories) are equally likely and for *n* = 5 there are *n*!(*n* − 1)!*/*2^*n*−1^ = 180 ranked topologies (Degnan and Salter 2005). Figure 5A shows that all 180 ranked topologies are sampled approximately uniformly as expected. Figure 5B shows that an individual terminal branch follows the expected distribution given in Equation 8 and Figure 5C shows that the sum of branch lengths in the sampled trees follows the expected distribution given in Equation 7.

**Figure 5:**
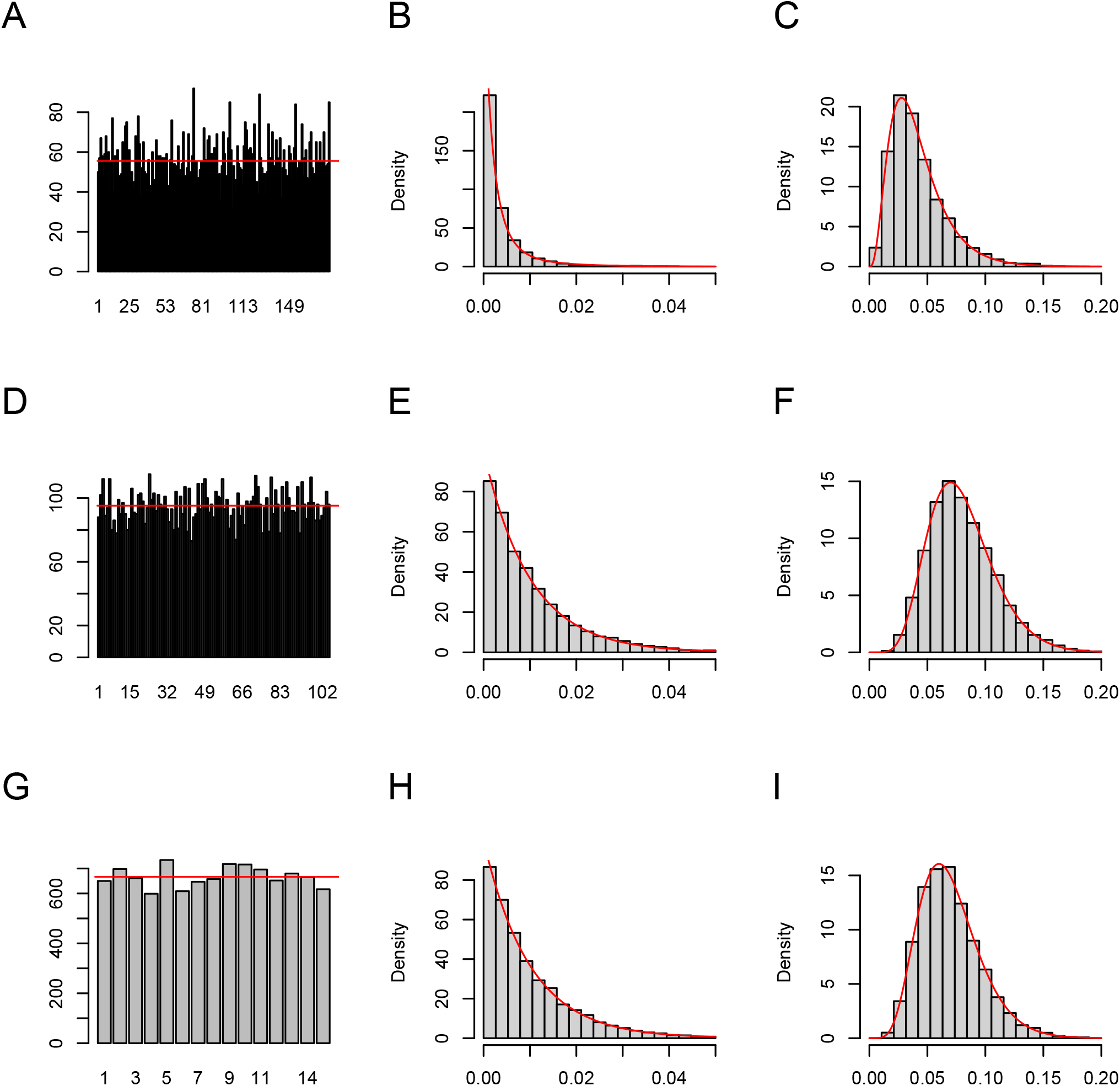
Sampled (histograms) and expected (lines) distributions under the ultrametric tree model (A-C), rooted model (D-F) and unrooted model (G-I) for the topology (A, D, G), for a single branch length (B, E, H) and for the sum of branch lengths (C, F, I).

Under the rooted tree model (Equation 9) all topologies are equally likely by definition and for *n* = 5 there are (2*n* − 3)!! = 105 of them. Figure 5D shows that all of them are sampled approximately uniformly as expected. Furthermore, Figures 5E and 5F show that terminal branch lengths and the sum of branch lengths in sampled trees follow the correct distributions from Equation 10, respectively. Finally, under the unrooted tree model (Equation 11) all topologies are also equally likely and for *n* = 5 there are (2*n* − 5)!! = 15 of them. Figure 5G shows that all of them are sampled approximately the same number of times as expected. Figures 5H and 5I show that terminal branch lengths and the sum of branch lengths follow the distributions from Equation 12 expected under the unrooted tree model. For each of the three tree types, we use a chi-square test to confirm that the sampled topologies (Figures 5A, D and G) are uniform as expected and we use an Anderson-Darling simple hypothesis test to confirm that the sampled terminal branch lengths (Figures 5B, E and H) and sums of branch lengths (Figures 5C, F and I) all follow their corresponding expected distributions.

These results provide numerical validation of the formulas for redundancy and Jacobians of each of the three models (Table 1). They also demonstrate that it is possible to combine a real-valued sampler with the link function in Equation 13 to sample from the desired phylogenetic distributions in Equations 6, 9 and 11. This includes in particular the ability to explore the tree space over different topologies despite the Jacobian calculations applying only within each topology. Finally, the possibility to sample phylogenetic trees using our parametrisations suggests that they could be used within a Bayesian phylogenetic inferential framework.

### 8 Toy example of Bayesian phylogenetic inference

We now show a small toy example of how our new unconstrained Euclidean parametrisations can be used to perform Bayesian phylogenetic inference, which aims to reconstruct a tree given some genetic data using Bayesian statistics to capture the posterior distribution of the tree (Nascimento et al. 2017). After observing data *D* made of *n* aligned genetic sequences of length *L*, the distribution of phylogenetic trees becomes the posterior with density

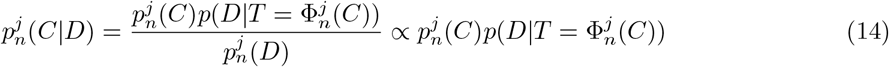

The first term in this product is the prior density from Equation 13 and the second term is the likelihood of the data given the phylogenetic tree 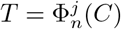 obtained using Algorithm 1. This likelihood can be calculated efficiently using the Felsenstein pruning algorithm (Felsenstein 1981) which is used in most phylogenetic inference tools.

For this toy example we focus on the ultrametric model as it is the simplest with just *n* − 1 parameters (Table 1). We generate a tree from the coalescent model (Equation 6) with *n* = 10 leaves and scaling factor *θ* = 0.005. This tree is shown in Figure 6A. A sequence of length *L* = 10, 000 bp is generated for the root of this tree by picking each of the four nucleotides {A,C,G,T} uniformly at random for each position. A sequence is then generated for each of the 10 leaves by evolving the sequence along the branches of the tree. Many substitution models can be used for this purpose (Nascimento et al. 2017) and here we use the simplest Jukes-Cantor model (Jukes and Cantor 1969) which avoids having to infer additional parameters since it considers all substitutions from any nucleotide to any other to be equally likely. The dataset *D* consists of the 10 sequences obtained for the leaves of the tree.

**Figure 6:**
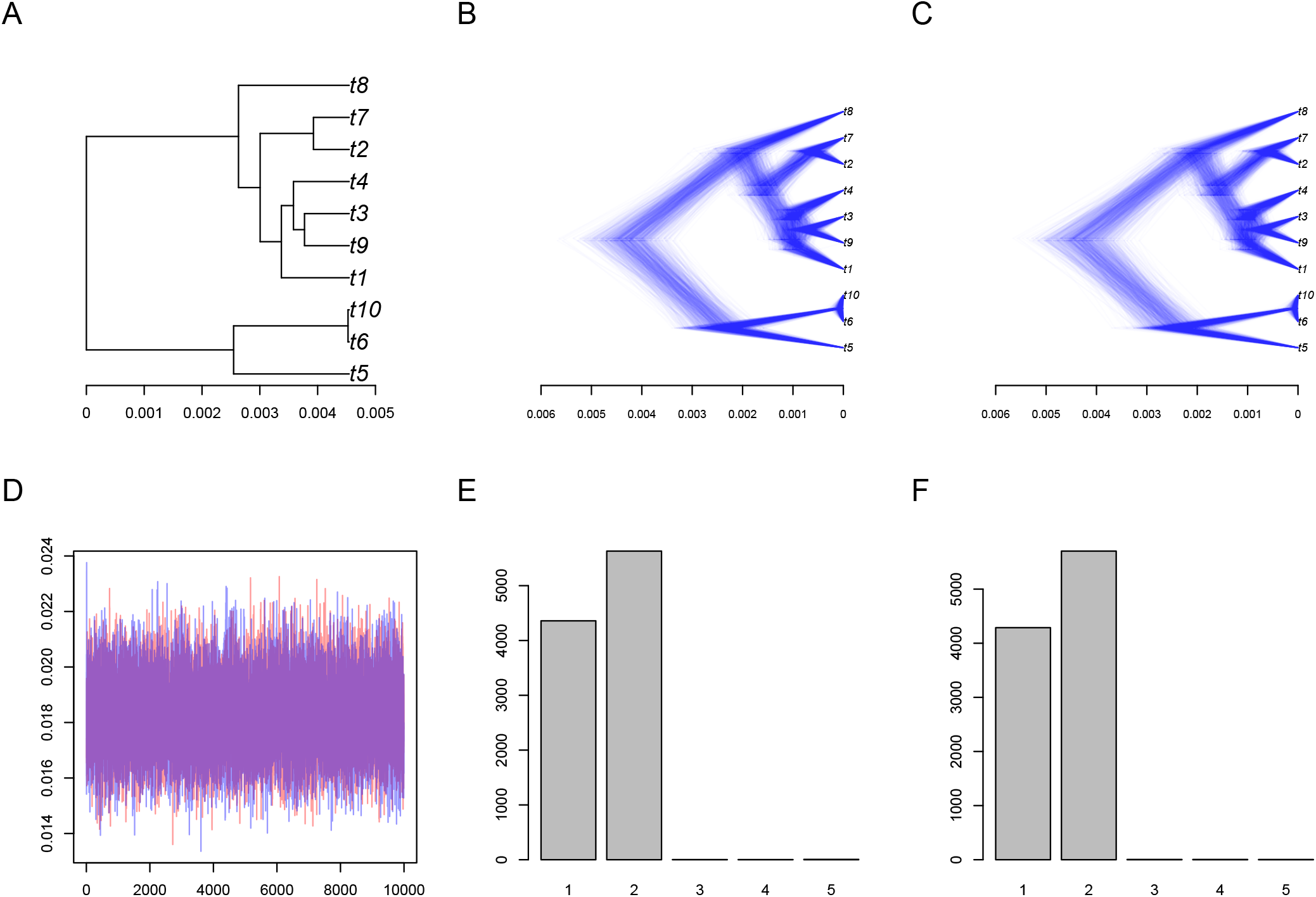
Toy example of Bayesian phylogenetic inference using two independent runs. (A) Correct tree used in simulation. (B-C) DensiTree representations of trees sampled in the two runs. (D) Traces of the sum of branch lengths in both runs. (E-F) Posterior count of topologies according to the two runs.

We perform inference using the same adaptive Metropolis sampler as described in the previous section, with the parameter being the *n* − 1 = 9 values in the real-valued vector *C*, and the target being the posterior distribution in Equation 14 given the data *D*. The sampler was run for 2 × 10^6^ iterations, with the first half discarded as burn-in and the remainder sampled every 100 iterations to produce a sample of size 10,000. Several runs are performed independently, and they are found to have converged and mixed well based on a Gelman-Rubin statistic below 1.1 and effective sample sizes above 1000 for the sum of the branch lengths and each terminal branch length (Plummer et al. 2006). The tree topology is also found in each run to have an approximate effective sample size greater than 1000 (Warren et al. 2017). Figures 6B and C show the phylogenetic posterior samples for two independent runs using a DensiTree graphical representation (Bouckaert 2010). The two posterior samples look very similar, and give high support to the correct phylogeny shown in Figure 6A. Figure 6D shows the trace of the two runs for the sum of branch lengths, again showing good mixing and agreement.

Finally, the Figures 6E and F show the posterior probability given to different topologies, with good agreement between them. The highest posterior probability was given to the correct topology, but significant weight was given to other similar topologies, showing that uncertainty remains that is being explored. Indeed the value of the scaling factor *θ* = 0.005 was selected to create a balanced situation, with enough polymorphism to inform the tree but not so much as to remove all uncertainties in the inference. Reducing this scaling factor decreases the number of substitutions so that the posterior becomes more similar to the prior without data (similar to Figure 5), whereas increasing the scaling factor adds more substitutions so that only a single topology becomes probable. The balanced situation shown in Figure 6 demonstrates our ability to explore the tree space, including different topologies when the posterior requires it.

## 9 Implementation

We implemented the methods described in this paper in a new R package entitled ContPhylo which is available at https://github.com/xavierdidelot/ContPhylo for R version 3.5 or later. This package contains all the code needed to reproduce all the results shown in this article. In particular, we implemented the Algorithms 1, 2 and 3 to construct phylogenetic trees from their Euclidean representations. We also implemented the reverse algorithms, which take as input a phylogenetic tree from one of the three types and outputs a canonical representation of this tree in the Euclidean space. In the canonical representation, children of every internal node are ordered by smallest descendant leaf label, so every new leaf is a second child: each graft coordinate is the larger of its two equivalent values given the preceding coordinates. We used ape and phangorn to store, manipulate and visualise phylogenetic trees (Paradis and Schliep 2019; Schliep 2011). We used the adaptive Metropolis sampler with coerced acceptance rate (Vihola 2012) implemented adaptMCMC (Scheidegger 2012). We used coda (Plummer et al. 2006) to analyse the MCMC outputs, and rwty (Warren et al. 2017) to compute the approximate effective sample size of tree topologies.

## 10 Discussion

The three constructions we introduced provide unconstrained Euclidean parametrisations of the three most commonly used types of phylogenetic tree spaces. The rooted ultrametric construction is the simplest case (Algorithm 1 and Figure 2). Each added leaf requires only its graft coordinate, because ultrametricity uniquely determines the new terminal-branch length. Allowing unrestricted branch lengths introduces one additional log-length parameter *b*_*i*_ at every grafting step (Algorithm 2 and Figure 3). Exponentiation of these parameters contributes the factors exp(*b*_*i*_) to the Jacobian (Equation 4). The unrooted construction applies the unrestricted rooted construction to the first *n* − 1 leaves, attaches the last leaf *n* to the root with length exp(*c*), and then forgets the root designation (Algorithm 3 and Figure 4). It therefore removes one graft coordinate relative to the rooted case and has the minimal dimension 2*n* − 3, with similar Jacobian as in the rooted case (Equation 5).

In every case, the dimension of the parameter space equals the number of parameters needed to specify the branch lengths for a given binary topology. All three functions are surjective onto the corresponding space of binary trees with strictly positive branch lengths and extend continuously to the completed tree space obtained by allowing zero-length branches. Their parameter spaces are divided into subspaces on which the selected contour branches and traversal directions are fixed. Within each such subspace, the resulting topology is fixed and the function is continuous. The boundaries between subspaces correspond to grafting at tree nodes or contour endpoints. At these boundaries a branch contracts to zero, so adjacent binary topologies meet at the same tree in the completed space, with the same order of the children owing to the convention on the order of the children of the new node, so that subsequent contour traversals also agree. The functions are therefore continuous globally.

In all three constructions, the redundancy of representations arises from the redundancy in the contour traversal (Figure 1). This redundancy is clearly a drawback of the parametrisations we proposed, but it seems unlikely that a bijection exists between phylogenetic tree space and Euclidean space. Since BHV space is not a manifold (Billera et al. 2001), no homeomorphism with ℝ^*d*^ exists. Restricting our parametrisation to canonical representatives (see Section 9) gives a bijection almost everywhere, but onto a tree-dependent subset of R^*d*^. At least the number of representations is fixed for each type of tree and value of *n* (Table 1) which allows us to sample without theoretical difficulty using Equation 13. In practice the multi-modality could be a problem for some samplers, although the different modes might be separate enough for only a single one to be explored. All modes are exact copies, so mode-hopping is harmless in terms of the sampled phylogenetic trees. This problem is therefore analogous to label switching in mixture and Hidden Markov Models (Papastamoulis 2016).

Many promising Monte-Carlo methods have not been applied in Bayesian phylogenetics because they can only be applied to continuous parameters. Our new unconstrained Euclidean parametrisation of phylogenetic trees provides an opportunity to use MCMC methods that had not previously been possible to use in Bayesian phylogenetics. As an example of this we used an adaptive Metropolis algorithm (Vihola 2011) to infer the phylogeny, and showed that it can sample the prior when no data is available (Figure 5) and the posterior when data is provided (Figure 6). This toy example of Bayesian phylogenetic inference was aimed as a proof-of-concept, not as a practical approach compared to current state-of-the-art methods such as MrBayes (Ronquist et al. 2012) or BEAST (Baele et al. 2025). The focus here was on providing constructions that enable the use of new algorithms, not on the application of these algorithms or on scalability to real-life data analysis. Our implementation uses R and is not optimised. More importantly, the adaptive Metropolis algorithm (Vihola 2011) we used is not the most promising Bayesian inference method unlocked by our new parametrisations. Future work should investigate the application of Hamiltonian Monte-Carlo (Betancourt 2017; Carpenter et al. 2017), Automatic Differentiation Variational Inference (Kucukelbir et al. 2017; Blei et al. 2017) or normalising flows (Papamakarios et al. 2021) to our parametrisations in order to develop new approaches to Bayesian phylogenetic inference.

